# *Let-7* miRNAs control auditory sensory progenitor behavior in the vertebrate inner ear

**DOI:** 10.1101/655993

**Authors:** Lale Evsen, Shuran Zhang, Angelika Doetzlhofer

## Abstract

The evolutionary conserved lethal-7 (*let-7*) family of microRNAs (miRNAs) is a well-known activator of terminal mitosis and differentiation. Surprisingly, we previously found that overexpression of *let-7* miRNAs in the murine auditory organ accelerated the terminal mitosis of auditory sensory progenitors (pro-sensory cells) but failed to stimulate their differentiation into mechano-sensory hair cells (HCs). To further address the role of *let-7* miRNAs in auditory sensory differentiation, we conducted gain and loss of function experiments in the developing chicken auditory organ, the basilar papilla (BP). Using a sponge approach, we show that the disruption of *let-7* miRNA function in the developing BP delays pro-sensory cell exit and delays differentiation of auditory HCs, revealing that endogenous *let-7* miRNAs limit pro-sensory cell self-renewal in the developing BP. However, consistent with the role of *let-7* miRNAs in the murine auditory organ, *let-7b* overexpression in the developing BP delayed HC differentiation, suggesting that too low or too high *let-7* miRNA levels disrupt HC differentiation. Furthermore, we provide evidence that the repressive role of *let-7* miRNAs in HC differentiation may be due to its targeting of the chromatin remodeler CHD7. Mutation in the human *CHD7* gene causes CHARGE syndrome, which amongst others is characterized by inner ear and hearing deficits. Using target prediction algorithms, we uncovered a highly predictive and evolutionary conserved *let-7* binding site within the *Chd7* transcript. Consistent with being a target of *let-7* repression, we demonstrate that *let-7b* overexpression significantly reduced CHD7 protein expression in to the developing BP. Furthermore, utilizing an inducible *let-7g* transgenic mouse model, we show that *let-7* miRNAs negatively regulate CHD7 protein expression in developing murine cochlear, retinal and brain tissue. CHD7 is dosage dependent and the here described regulation by *let-7* miRNAs may be critical to fine tune CHD7 protein levels during sensory and neuronal development.

**SIGNIFICANCE:** The evolutionary highly conserved *let-7* miRNAs are essential for proper timing of cell state transitions during embryogenesis. Even though abundantly expressed in the vertebrate auditory organ, surprisingly little is known about their function in auditory sensory differentiation. Here, we demonstrate that endogenous *let-7* miRNAs are essential for limiting auditory sensory progenitor (pro-sensory) cell self-renewal. Furthermore, we find that precocious *let-7* miRNAs expression interferes with auditory hair cell differentiation and identify chromatin remodeler CHD7 as a potential target gene of *let-7* repressive function in HC differentiation.

## INTRODUCTION

The vertebrate auditory organ contains an array of highly specialized mechano-sensory cells-called hair cells (HCs) that are critical for our ability to detect sound. Auditory HCs and their surrounding supporting cells (SCs) derive from a common pool of progenitor cells-referred to as pro-sensory cells (Fekete et al., 1998). SOX2, a high-mobility group transcription factor is a master regulator of inner ear sensory development. Early otic ablation of SOX2 abolishes the formation of both vestibular and auditory pro-sensory cells (Kiernan et al., 2005; Neves et al., 2011; Pan et al., 2013). In addition, SOX2 is required for transcriptional activation of *Atoh1* expression in the developing inner ear (Ahmed et al., 2012; Kempfle et al., 2016; Neves et al., 2012). ATOH1, a bHLH transcription factor, is the earliest marker of nascent HCs and is both necessary and sufficient for the formation of auditory and vestibular HCs (Bermingham et al., 1999; Chen et al., 2002; Zheng and Gao, 2000).

In the murine cochlea, terminal mitosis and differentiation of auditory pro-sensory cells follow steep longitudinal gradients that initiate at opposing ends of the sensory epithelium. The timing of these seemingly uncoupled events is controlled, amongst others, by the *let-7* family of microRNAs (miRNAs) and its antagonist the RNA-binding protein (RBP) LIN28B (Golden et al., 2015). *Let-7* (lethal-7) and *LIN-28* were initially discovered in *C. elegans*, being part as members of a cascade of genes that regulate the timing of developmental events (heterochrony)(Ambros and Horvitz, 1984; Reinhart et al., 2000). Vertebrates have multiple let-7 isoforms (species) and to date, nine mature *let-7* miRNA species (*let-7a, let-7b, let-7c, let-7d, let-7e, let-7f, let-7g, let-7i, mir-98*) have been identified in humans and mice (reviewed in (Roush and Slack, 2008). As for most miRNAs, the primary transcript of *let-7* miRNA species is processed by the RNase III enzymes, Drosha and Dicer into ∼ 22 nucleotide miRNA duplex, of which one strand termed the lead strand is incorporated into the miRNA-induced silencing effector complex (miRISC) (reviewed in (Yates et al., 2013)). *Let-7* miRNA isoforms share a conserved seed sequence, which is used to recognize and to bind to target mRNAs. Recruitment of miRISC complex to the *let-7* target mRNA results in deadenylation, leading to mRNA decay or translational repression (Wakiyama et al., 2007). *Let-7* miRNAs limit self-renewal and promote a terminal differentiated by targeting pro-growth genes such as *Lin28a, Lin28b, Hmga2 and Igf2bp1 (Imp1)*. In turn, LIN28B and the closely related RBP LIN28A maintain stemness and proliferation by interfering in the processing of primary *let-7* transcripts into functional mature miRNAs (Nishino et al., 2013; Piskounova et al., 2008; Viswanathan et al., 2008; West et al., 2009).

Similar to other neuronal and sensory tissues, mature *let-7* miRNA expression in the developing cochlea steadily increases during terminal differentiation. However, surprisingly, *let-7g* overexpression in the murine cochlea failed to trigger premature HC differentiation. Instead, we found that *let-7g* overexpression mildly delayed HC differentiation (Golden et al., 2015). To further characterize the function of *let-7* miRNAs in auditory sensory development, we manipulated *let-7* levels in the avian auditory organ, the so-called basilar papilla using *in-ovo* electroporation. We found that disruption of *let-7* function in the developing chicken BP prolongs pro-sensory cell proliferation and delays HC differentiation, indicating that endogenous *let-7* miRNAs play a key role in restricting pro-sensory cell self-renewal. Furthermore, we uncovered that increased *let-7* expression inhibits both sensory progenitor proliferation and HC differentiation, suggesting that pro-sensory expression of mature *let-7* miRNA species have to be precisely fine-tuned to ensure proper timing of terminal mitosis and differentiation. Finally, we identified CHD7, a key regulator of sensory-neural development, as a novel target of *let-7* miRNAs. We demonstrate that the CHD7 gene harbors a highly predictive and evolutionary conserved *let-7* binding site and demonstrate that in chicken and mice overexpression of *let-7* miRNAs significantly reduces CHD7 protein expression within auditory sensory epithelium as well as retinal and brain tissues.

## MATERIALS AND METHODS

### Eggs, *in ovo* micro-electroporation, and expression constructs

Fertilized chicken eggs (B&E York Springs, PA) were incubated at 37° C and staged according to Hamburger and Hamilton (Hamburger and Hamilton, 1951) at HH24-25 (E4-E4.5). To over-express *let-7* we electroporated a dual promoter lentiviral plasmid let-7b-GFP, in which the murine primary-*let-7b* sequence (NR_029727.1) is expressed under the control of the mouse U6 promoter and GFP expression is driven by a CMV promoter. Let-7b-GFP plasmid was a gift from Sean Morrison (Addgene plasmid # 25404) (Nishino et al., 2008). To knock-down *let-7* function, we used the “*Let7-sponge*” construct, which contains six canonical *let-7-5p* binding sites cloned downstream of *U6* promoter. pRNA-U6-let7 sponge was a gift from Phillip Zamore (Addgene plasmid # 35664). To visualize electroporated cells we co-electroporated *pMES-IRES-GFP* in which the expression of *GFP* is driven by a chicken *β-actin* promoter. Plasmids were delivered to the right otic vesicle targeting the BP rudiment at E4-E4.5 (HH24-25) by electroporation as previously described (Evsen and Doetzlhofer, 2016) with the left ears serving as untreated internal controls. Briefly, this was conducted by filling the right otic vesicle with plasmids at a concentration of 3-5 μg/μl tinted with 0.1 % Fast Green. A negative 2 mm platinum electrode was placed anterior-ventral to the right otic vesicle and the positive electrode placed in parallel 1 mm apart. Four pulses at 12 volts with 100 ms duration and 200 ms spacing were applied using an ECM 830 square wave electroporation system (BTX Harvard Apparatus). Then, the eggs were sealed and returned to the incubator for 6, and 18 to 44 hrs.

### Mice breeding and genotyping

All experiments and procedures were approved by the Johns Hopkins University Institutional Animal Care and Use Committees protocol, and all experiments and procedures adhered to National Institutes of Health-approved standards. iLet-7g transgenic mouse model: transgenic mice that express degradation resistant form of *let-7g* under the control of a tetracycline response element were mated with transgenic mice that ubiquitous expressed modified reverse tetracycline-trans activator (*M2-rtTA*). The *let-7*g transgenic mice were obtained from George Q. Daley (Children’s Hospital, Boston, MA) (Zhu et al., 2011). The *M2-rtTA* mice were purchased from Jackson Laboratories (stock no. 006965). Mice were genotyped by PCR using the following primers. iLet-7: (ColfrtA/ GCA CAG CAT TGC GGA CAT GC, ColfrtB/ CCC TCC ATG TGT GAC CAA GG, ColfrtC/ GCA GAA GCG CGG CCG TCT GG). M2-rtTA: (rtTAA/ GCG AAG AGT TTG TCC TCA ACC, rtTAB/ AAA GTC GCT CTG AGT TGT TAT, rtTAC/ GGA GCG GGA GAA ATG GAT ATG). Mice of both sexes were used in this study. All mouse lines were maintained on a mixed background of C57BL/6 and CD-1. Embryonic development was considered as E0.5 on the day a mating plug was observed. To induce *let-7g* expression, dox was delivered to time-mated females via ad libitum access to feed containing 2 g of doxycycline per kg of feed (Bioserv no. F3893), which was continued until the stage the tissue was harvested. Dox treatment was begun at E10.5 and embryos were removed at E11.5 and E13.5 from timed-mated females and staged by using the EMAP eMouse Atlas Project Theiler staging criteria (www.emouseatlas.org/emap/ema/home.html).

### RNA *in situ* hybridization, immunohistochemistry, and *in ovo* EdU incorporation assay

Chicken and mouse embryos were staged using Hamburger Hamilton staging criteria (Hamburger and Hamilton, 1951) and EMAP eMouse Atlas Project (http://www.emouseatalas.org) Theiler staging criteria respectively. Whole embryos or heads were briefly fixed in 4% (vol/vol) paraformaldehyde (PFA) (Electron Microscopy Sciences) in PBS, cryoprotected using 30% sucrose in PBS, and embedded in OCT (Sakura Finetek). Tissue was sectioned at a thickness of 14 µm and collected on SuperFrost Plus slides (Thermo Scientific) and stored at −80°C. Tissue was sectioned at a thickness of 14 µm and collected on SuperFrost Plus slides (Thermo Scientific) and stored at −80°C. *In situ* hybridization assays to detect mRNA and mature miRNA transcripts were carried out as previously described (Raft et al., 2007) (Golden et al., 2015). Digoxigenin (Dig)-labeled anti-sense RNA probes were used to detect *Gfp* and chicken *Atoh1, Lfng* and *Sox2* mRNA. Locked nucleic acid (LAN) probes (Exiqon) were used for *in situ* detection (Obernosterer et al., 2007) of mature *let-7g-5p, let-7b-5p, let-7c-5p*, and *mir-183-5p* miRNAs. The primary antibodies used were goat polyclonal α-GFP-(FITC) (1:400; GeneTex), rabbit polyclonal α-SOX2 (1:4000 with 10 min antigen retrieval in citrate buffer; Chemicon), goat polyclonal α-SOX2 (1:500; Santa Cruz Biotechnology), goat polyclonal α-JAG1 (1:500; Santa Cruz Biotechnology), rabbit polyclonal α-CHD7 (1:500; Cell Signaling), rabbit polyclonal α-activated Caspase3 (1:800 with 10 min antigen retrieval in citrate buffer; Cell Signaling), mouse monoclonal α-phospho Histone 3 (1:500; Millipore), mouse monoclonal α-Tuj1-IgG2a (1:500; BioLegend), rabbit polyclonal α-MyosinVIIa (Myo7a) (1:1000 Proteus Biosciences), and mouse monoclonal α-Hair Cell Antigen (HCA; 1:1000, gift from Guy Richardson). The secondary antibodies were goat α-rabbit Alexa Fluor 488 (1:250; Invitrogen), goat α-mouse IgG2a 647 (1:250; Sigma), donkey α-goat Alexa Fluor 546 (1:250; Invitrogen), donkey α-rabbit Alexa Fluor 546 (1:250 Invitrogen), donkey α-mouse Alexa Fluor 647 (1:250; Invitrogen), and donkey α-rabbit Alexa Fluor 647 (1:250; Invitrogen). Antibody labeling was performed according to manufacturer’s recommendations. Immuno-labeled tissue sections were counterstained with Hoechst stain (blue). Image J software was used to quantify relative fluorescent intensity. For analysis of cellular proliferation in the developing chicken, 50 µl of 0.250 mg/ml EdU (5-ethynyl-2’-deoxyuridine) in 1x PBS sterile-filtered was added to the whole embryo *in ovo* as previously described (Evsen and Doetzlhofer, 2016). Click-iT AlexaFluor-488 or −546 Kit (Invitrogen) was used to detect incorporated EdU according to the manufacturer’s specifications.

### Western blots

Whole brain tissue obtained from control and ilet-7 transgenic embryos stage E11.5 was lyzed in RIPA lysis buffer (Sigma) supplemented with Roche Protease Inhibitor (Sigma) and Phosphatase Inhibitor Cocktail no.2 and no.3 (Sigma). Following manufactures recommendations, equal amounts of otic vesicles or whole brain protein extract were resolved on NuPAGE 4-12% Bis-Tris Gels (Invitrogen) or NuPAGE 3-8% Tris-Acetate Gels (Invitrogen) and transferred to Immun-Blot PVDF membrane (Bio-rad) by electrophoresis. Membranes were blocked in 5% no-fat dry milk in TBST and immunoblotted with rabbit polyclonal anti-CHD7 1:2000 (Cell Signaling Cat. No.6505S), rabbit polyclonal anti-SOX2 1:2000 (Millipore Cat. No. AB5603), rabbit polyclonal anti-TRIM71 1:2000 (gift from Gregory Wulczyn), goat anti-JAG1 1:400 (Santa Cruz Cat. No. sc-6011), and mouse α-β-actin HRP 1:400 (Santa Cruz, no.sc-47778 HRP) used as loading control. The HRP-conjugated secondary antibodies used are goat anti-rabbit IgG 1: 5,000 (Jackson ImmunoResearch no.111-035-003) and donkey anti-goat IgG (H+L) 1: 2,000 (Life Technologies Cat. No. A16005). Signal was revealed using either SuperSignal West Pico PLUS Chemiluminescent Substrate (Thermo Scientific) or SuperSignal West Femto Maximum Sensitivity Substrate (Thermo Scientific) according to manufacturer’s instructions. The intensity of each band was determined by densitometry, using ImageJ software. Protein levels were normalized using beta-actin.

### RNA extraction and qPCR

For RNA extraction, chicken auditory epithelia were isolated from wild-type chicken BPs by incubating the tissue in Thermolysin (500 μg/ml in 1X PBS, Sigma) for 20 min at 37**°** C. For each experiment a minimum of four auditory epithelia were pooled per condition and miRNeasy Micro kit (QIAGEN) was used to isolate total RNA and mRNA was reverse transcribed into cDNA using iSuperscript kit (Bio-Rad). QPCR was performed using SYBR Green kit (Life Technologies) and gene specific primer sets on a StepOne Plus PCR Detection System (Applied Biosystems Life Technologies). Each qPCR reaction was carried out in triplicate. Relative gene expression was analyzed using the ΔΔCT method (Livak and Schmittgen, 2001). The ribosomal protein L19 gene *RPL19* was used as endogenous reference gene. The following primers were used for qPCR in the chicken BP: RPL19-F CAG GAA GTT AAT TAA GGA TGG TTT GA; RPL19-R CAT CGT GCC CGA GAG TGAA; Atoh1-F CAA CGA CAA GAA GCT CTC CAA GT; Atoh1-R GGG CGC TGA TGT AGA TTT GC; Trim71-F GGC ACT CTG GAA GCA CTT TGA; Trim71-R AAG TGG CCC TCG TGA TTG AA; Nefm-F TCG AAA TTG CTG CAT ACA GGA A; Nefm-RCCA GAG AAG GCA CTG AAT CTT GT; Hmga2-F TGG CCT CAA CAA GTG GTT CA; Hmga2-R TCT TGT GAC GAT GTT TCT TCA GTC T; Lin28b-F GCC TTG AAT CAA TAC GGG TAA CA; Lin28b-R GGG TCG TCT TTC ACT TCC TAA ACA.

To quantify the expression of *let-7a-5p, let-7a-3p, let-7b-5p, let-7c-5p, let-7c-1-3p, let-7g-5p, let-7i-5p*, and *mir-125b-5p* transcripts, predesigned TaqMan Assays (Applied Biosystems, Life Technologies) were used according to manufacturer’s instructions. The snoRNA U6 was used as an endogenous reference gene for TaqMan-based miRNA measurements.

### Cell counts

In situ and immuno-labeled tissue sections were imaged using bright-field/ epi-fluorescent and confocal microscopes (ZEISS). Images from experimental and corresponding control tissue sections were assembled and analyzed in Photoshop CS6 (Adobe) and LSM image viewer (Zeiss). The number of SOX2(+) pro-sensory cells and SOX2 (+) EdU(+) pro-sensory cell within BP epithelia were manually counted. 4-6 tissue sections spanning the apical-basal extent were analyzed per animal and treatment. Data shown is mean percentage of SOX2(+) EdU(+) pro-sensory cells per section. Similarly, the number of phospho-H3(+) or activated-Casp3(+) cells within otic or cochlear epithelia were manually counted. 5-6 tissue sections of E11.5 otic tissue spanning the dorsal-ventral extent and 4-6 tissue sections spanning the apical-basal extent were analyzed per animal and genotype. Data shown is the mean number of activated-Casp3 (+) cells/section or the mean number phospho-H3 (+) cells/ section.

### Statistical Analyses

Values are presented as mean ± standard error (SEM), n = animals per group. All results were confirmed by at least two independent experiments. Two-tailed Student’s t-tests were used to determine confidence interval. P-values ≤ 0.05 were considered significant. P-values > 0.05 were considered not significant.

## RESULTS

### Opposing cell-cycle exit and HC differentiation gradients in the chicken BP

In the murine cochlea pro-sensory cell cycle exit and differentiation occurs in opposing longitudinal gradients. Pro-sensory cells located at the apex are the first to withdraw from the cell cycle but are last to differentiate into HCs and SCs (Chen et al., 2002; Chen and Segil, 1999; Ruben, 1967). Interestingly, opposing gradients of cycle exit and differentiation have also been reported for the chicken BP. In the chicken BP pro-sensory cell-cycle exit has been reported to start at around ∼ E5 (HH26) near the proximal (basal) end (Katayama and Corwin, 1989), whereas stereocilia bearing HCs have been reported to first appear at the distal (apical) end of the sensory epithelium at around ∼E6.5 (HH29) (Cotanche and Sulik, 1984). A Subsequent study that used an antibody against 275 kD hair cell antigen (HCA) revealed that the early distal-to proximal gradient of HC differentiation quickly disappears and a narrow band of HCs is present throughout the BP and HCs are added radially (Goodyear and Richardson, 1997). To confirm the previously reported pattern of cell cycle exit and differentiation we added thymidine analog EdU to whole chicken embryos at E4.5, E5.5, and E6.5, and analyzed EdU incorporation and HC differentiation one day later (Fig. 1A, and Fig. S1). Addition of EdU at E4.5 labeled a large fraction of SOX2(+) pro-sensory cells in the apex, indicating that at E4.5 the majority of pro-sensory cells located in the apex of the BP are mitotic. In contrast, at the base of the BP only few SOX2(+) pro-sensory cells, located at the abneural edge of the pro-sensory domain were labeled by EdU, suggesting that by E4.5 many of pro-sensory cells located in the base of the BP have already withdrawn from the cell cycle. A similar pattern of EdU incorporation was observed when EdU was added at E5.5 (Fig. S1, top panel). When EdU was added at E6.5 and analyzed at E7.5 only a few remaining EdU(+) cells were present on the neural edge of the differentiated BP sensory epithelium (Fig. S1, bottom panel).

**Figure 1.**
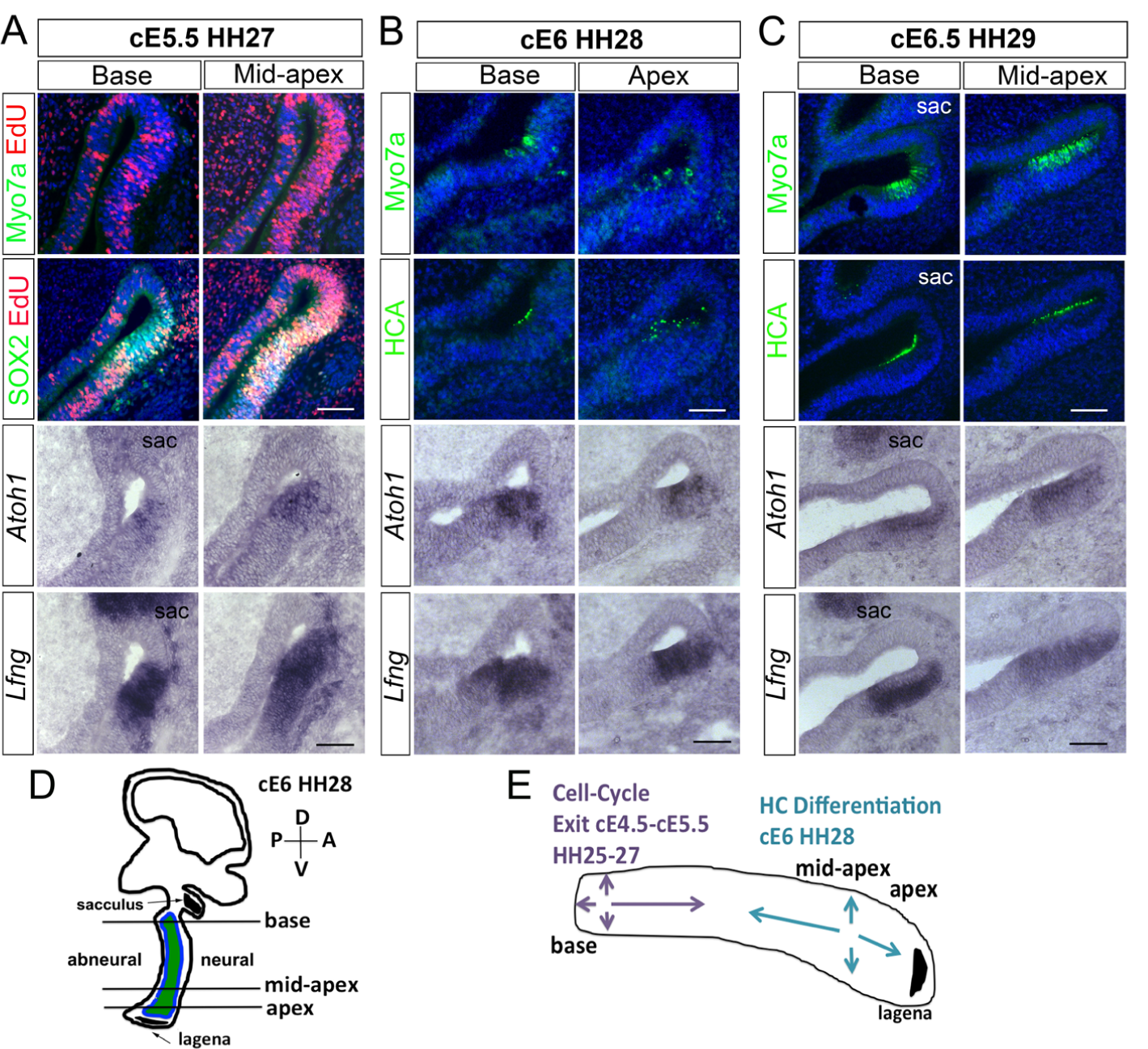
Characterization of terminal mitosis and hair cell (HC) differentiation in the developing chicken BP. **(A-C)** Shown are adjacent tissue-sections through the base (proximal) and apex (distal) segment of wild-type BPs stages (E5.5-E6.5; HH27-29). Myo7a and HCA immuno-staining (green) marks HCs, *Atoh1* ISH marks HC precursors and HCs, *Lfng* ISH and SOX2 immuno-staining marks the pro-sensory/sensory domain within the developing BP. EdU was added at E4.5 and incorporation was analyzed at E5.5. **(A)** Pro-sensory cells (SOX2+, *Lfng+*) located at the base incorporated EdU at a lower rate than pro-sensory cells located more apically. At E5.5 no Myo7a(+) HCs have yet formed but *Atoh1*(+) HC precursors are present within the *Lfng*(+) pro-sensory domain with the highest number in the mid-apex. **(B)** At stage E6.5, the number of *Atoh1*(+) HC precursors/HCs increases and the sensory epithelium contains few scattered Myo7a (+) and HCA(+) HCs cells. **(C)** At E6.5 the number of Myo7a(+) HCA(+) HCs increases and the majority of *Atoh1*(+) cells reside within the HC layer and *Lfng* expression is limited to the SC layer. Abbreviations sac, sacculus. **(D)** Schematic of the chicken inner ear ∼stage E6.0 indicating the position and orientation of shown tissue sections. Note that the developing BP is flanked by the sacculus and the lagena. **(E)** Summary model of terminal mitosis and differentiation in the developing BP. Scale bars 100 µm.

Taken together our results confirm the previous reports that pro-sensory cell cycle exit in the BP is starting at ∼E4.5 (HH25) and progresses from the base to the apex and from the center to periphery (Fig. 1 E) (see (Katayama and Corwin, 1989)). Furthermore, consistent with previous reports, we found that HC differentiation, as judged by HC-specific Myo7a and HCA expression, begins just prior to E6 (HH28). At E5.5 (HH27) no Myo7a(+) HC were detected within the BP (Fig. 1 A). However, *Atoh1*(+) cells could be readily detected within the pro-sensory domain, marked by *Lfng*, with the highest number of *Atoh1*(+) cells in the mid-apex (Fig. 1A). Interestingly, not all of the *Atoh1(+)* cells were confined to the HC layer (top layer), suggesting that in the developing BP *Atoh1* expression marks both HC precursors as well as differentiating HCs. Furthermore, the *Atoh1*(+) HC precursors/HCs cells were located close to the neural edge of the *Lfng*(+) pro-sensory domain, suggesting that tall HCs located at the neural edge of the sensory epithelium differentiate prior to short HCs located at the abneural edge of the sensory epithelium. At E6.0 (HH28) the number of *Atoh1*(+) cells in the base was increased compared to E5.5 (Fig. 1 A, B) and we observed few, scattered Myo7a(+) and HCA(+) HCs throughout the BP, with the highest number of Myo7a(+) and HCA(+) HCs being located in the apex (Fig. 1 B). At E6.5 the number of Myo7a(+) and HCA(+) HCs further increased in both base and apex and *Atoh1* and *Lfng* expression were confined to the HC layer and SC layer respectively (Fig. 1 C).

### *Let-*7 miRNAs are highly expressed in differentiating HCs

To gain insights into the role of *let-7* miRNAs in chicken auditory development, we first analyzed *let-7* target gene (*Lin28b, Trim71* and *Hmga2*) and mature *let-7* miRNA expression in enzymatically purified chicken BPs stages E5.5 –E15.5 using RT-qPCR. Our analysis revealed that prior to the onset of HC differentiation (E5.5), *Lin28b, Trim71* and *Hmga2* were abundantly expressed, but their BP-specific expression quickly declined during the early (E6.5) and late phases (E7.5, E8.5) of HC differentiation (Fig. 2 A). Conversely, the lead strand (5-p) of mature *let-7* miRNA species (*let-7a, b, c, g, i)* and the functionally related *mir-125b* were 2-20-fold higher expressed in the terminal differentiated BP (E9.5) compared to the undifferentiated BP (E5.5). The expression of mature *let-7b, let-7c* and *let-7i* miRNAs further increased during HC maturation and were 2-3-fold higher expressed in the mature BP (E15.5) compared to immature terminal differentiated BP (E9.5) (Fig. 2 B). We also examined the expression of the *let-7a* and *let-7c* passenger strands (3-p) in developing chicken and murine auditory sensory epithelia. As expected, the passenger strands (3-p) were produced at much lower frequency (∼20-100 fold less) than the corresponding lead strands (5-p) (Fig. S 2).

**Figure 2.**
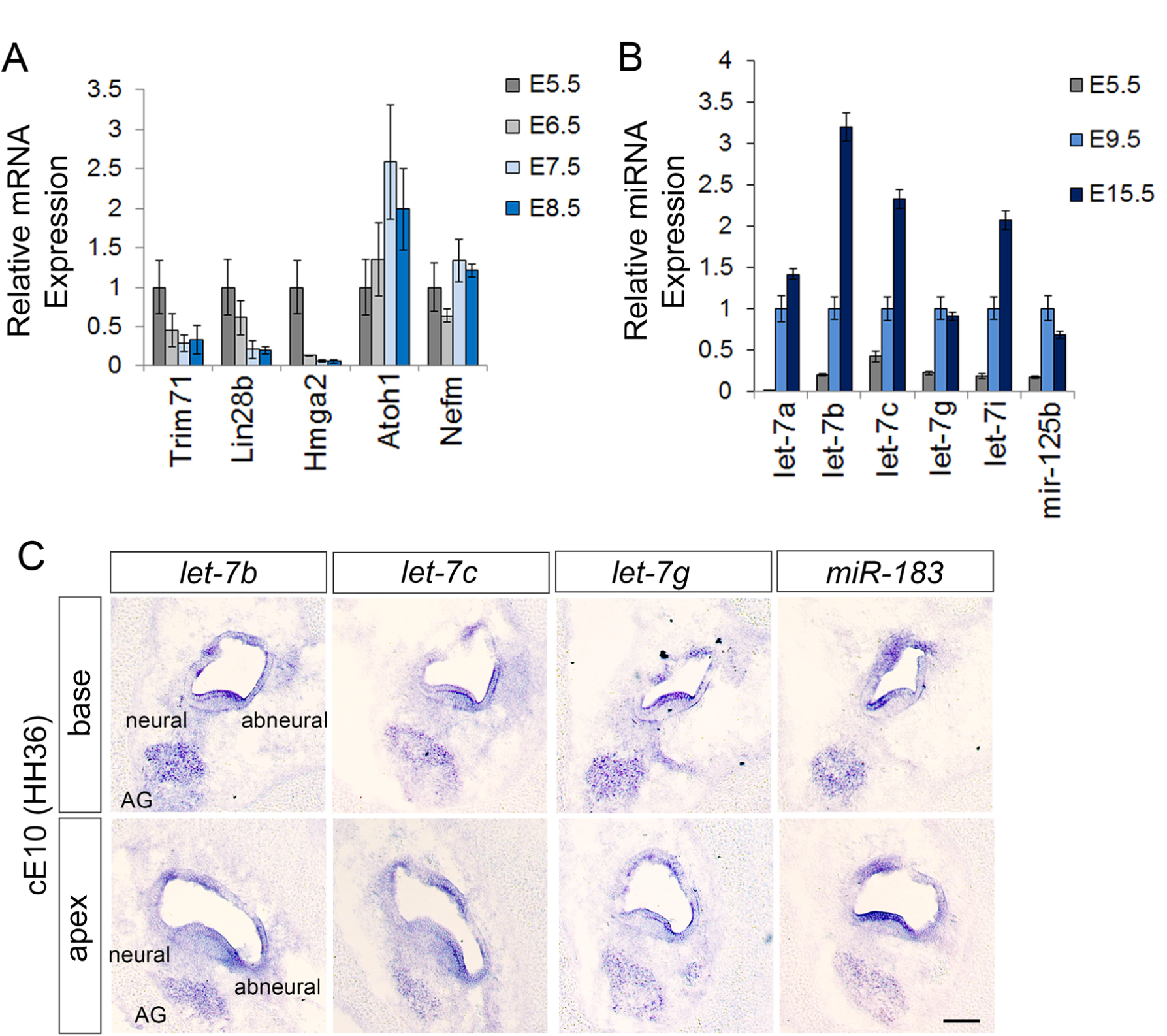
Spatial and temporal patterns of *let-7* miRNA expression in the developing chicken BP. Relative mRNA (A) and mature miRNA (B) transcript abundance was assayed by RT-qPCR in enzymatically purified wild-type BP epithelia. Cell-type specific expression of mature miRNAs was analyzed using LNA miRNA ISH assays. **(A)** The relative high mRNA expression of *let-7* target genes *Trim71, Lin28b*, and *Hmga2* at E5.5 declines in the differentiating BP (E6.5, E7.5 and E8.5), as *Atoh1* expression increases. Data are ± SEM (n = 3, technical replicate). **B)** *Let-7a-5p, let-7b-5p, let-7c-5p, let-7g-5p, let-7i-5p*, and *mir-125b-5p* miRNA expression steadily increases during BP differentiation (E5.5-E9.5) and continues to increase in the post-differentiated BP at E15.5. Data are ± SEM (n = 3, technical replicate). **(C)** *Let-7b-5p, let-7c-5p, let-7g-5p* and *miR-183-5p* miRNAs are highly expressed within the HC layer and acoustic ganglion (AG). Note, *let-7* miRNAs are expressed in a radial gradient within the auditory sensory epithelium, with higher expression in cells at the abneural side than neural side. Scale bar 100 µm.

Next, we used Locked Nucleic Acid nucleotides (LNAs)-based *in situ* hybridization (ISH) assay to characterize the spatial expression pattern of mature *let-7* miRNAs in the terminal differentiated chicken BP stage E10 (HH36) BP. As positive control we analyzed the expression of *mir-183-5p*, a miRNA previously shown to be selectively expressed in HCs and auditory ganglion neurons (Zhang et al., 2015). Our analysis revealed that within the BP *let-7b-5p, let-7c-5p*, and *let-7g-5p* are highest expressed in HCs and the auditory ganglion, showing partially overlapping expression with *miR-183-5p* (Fig. 2 C). Interestingly, the expression of *let-7* miRNAs in HCs differed along the basal-apical and neural-abneural axes, with most pronounced expression in abneural (short) HCs located at the basal (proximal) portion of the BP. This graded pattern of expression may have functional implication for the maturation and specialization of tall (neural) and short (abneural) HCs within the BP.

### Inhibition of endogenous *let-7* miRNAs increases pro-sensory cell proliferation and delays HC differentiation

In chicken the *let-7* family of miRNAs consists of at least 9 members (*let-7a, b, c, d, f, g, i, j, k*) (www.mirbase.org/). To disrupt the function of all *let-7* miRNA family members we overexpressed a *let-7 “sponge”* RNA that contains an array of canonical *let-7-5p* binding sites, complementary to the seed sequence shared by all *let-7* family members and across species. When overexpressed at sufficiently high levels the *let-7* sponge functions as a decoy for endogenous *let-7* miRNAs, sequestering *let-7* miRNAs from their endogenous target. We co-electroporated the *let-7 sponge* together with a *GFP* reporter in E4.5 chicken embryos to ensure proper targeting to the BP rudiment (Fig. S3). EdU was added at time of electroporation and specimens were analyzed 44 hours later. We found that *let-7sponge* over-expression in the developing BP led to a significant increase in the number of pro-sensory cells that incorporated EdU compared to left internal untreated BP (control) (Fig. 3 A, B), suggesting that in the developing BP *let-7* miRNA function in fine-tuning pro-sensory cell proliferation. Moreover, BPs that received the *let7-sponge* contained either no (n=3/9) or fewer Myo7a(+) HCs (n=6/9) in the apex to mid-apex than control (Fig. 3 C, D). Furthermore, we found that more than half (n=5/8) of *let7-sponge* overexpressing BPs had no or fewer *Atoh1*(*+*) cells in the apex and mid-apex compared to control (Fig. 3 C, D). To verify that electroporation by itself did not alter pro-sensory cell behavior, we compared pro-sensory proliferation and differentiation in mock electroporated BPs (GFP only) and left untreated BPs (control). Our analysis revealed no differences in pro-sensory cell proliferation or HC differentiation (Fig. S3). Taken together, our findings indicate that *let-7* miRNAs in the developing BP are required for limiting pro-sensory proliferation as well as for the timely induction of HC differentiation.

**Figure 3.**
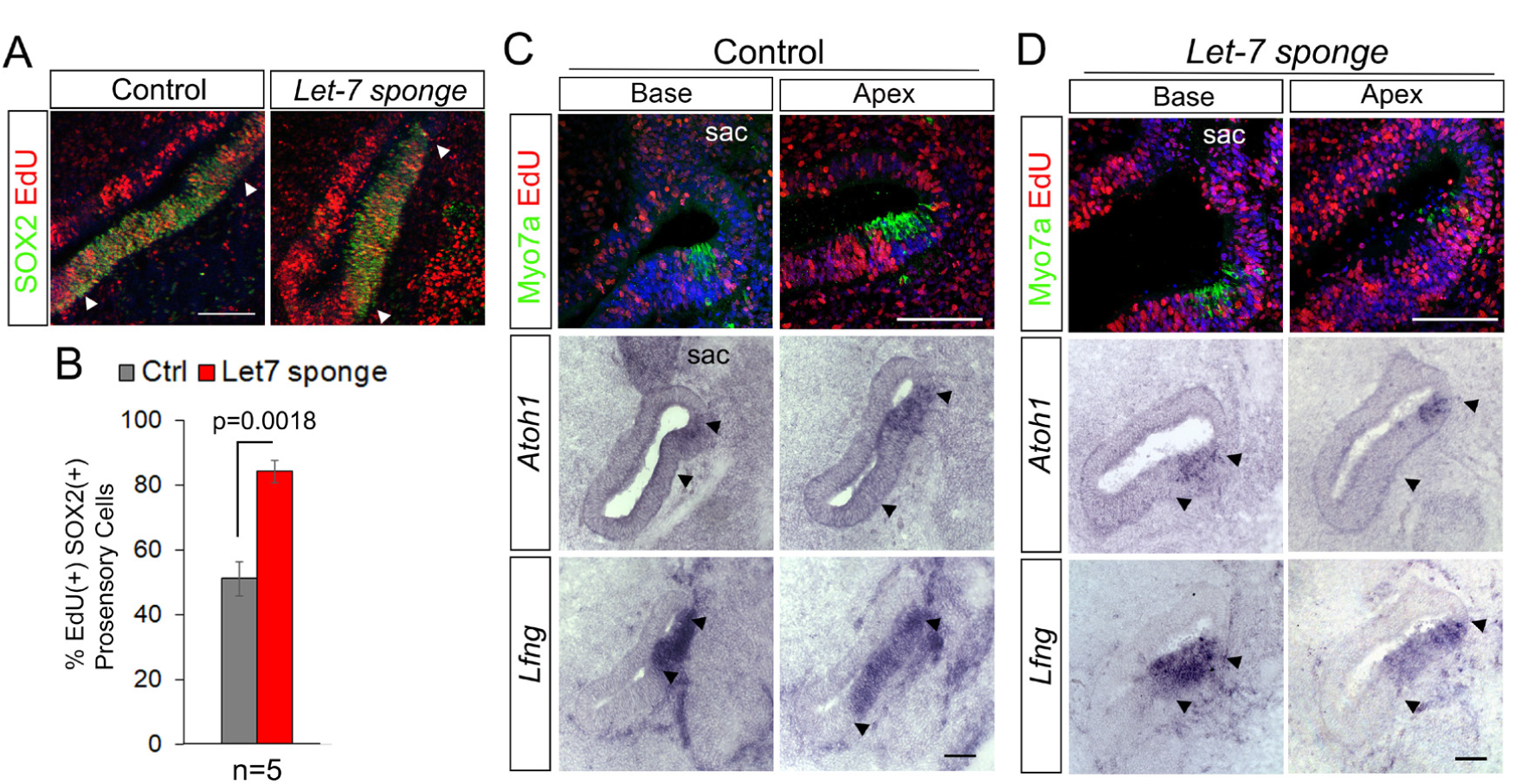
Disruption of *let-7* miRNA function increases pro-sensory cell proliferation and delays HC formation in the developing BP. Shown are adjacent sections through the base or apex of untreated control BPs (A, C) and of *let7-sponge* electroporated BPs (B, C) 44 hours after electroporation and EdU treatment at E4.5. Note white and black arrowheads mark pro-sensory domain. **(A-B): *Let7-sponge* treatment increases pro-sensory cell proliferation.** EdU incorporation in SOX2(+) pro-sensory cells is significantly increased in the mid-apex of *let-7* sponge treated BPs (A, left panel) compared to corresponding control BPs (A, right panel). **(B)** Quantification of A. Data are mean ± SEM (n = 5 animals per group, p=0.0018 by Student’s t-test, p ≤ 0.05 is deemed significant). **(C-D) *Let7-sponge* treatment delays HC differentiation.** The apex of the *let-7* sponge treated BP contains fewer Myo7a(+) HCs and fewer *Atoh1*(*+*) HC precursors (D) compared to the apex of control BPs (C). Note the size of the *Lfng*(+) pro-sensory/sensory domain, analyzed in adjacent sections, is unchanged in *let-7* sponge treated BPs compared to control. Abbreviations: sac, sacculus. Scale bars 100 µm.

### Overexpression of *let-7b* reduces pro-sensory cell proliferation and interferes with auditory HC differentiation

Next, we examined whether *let-7* miRNA overexpression is sufficient to trigger premature HC differentiation in the developing chicken BP. To increase the abundance of mature *let-*7 miRNAs, we electroporated E4.5 chicken embryos targeting the BP rudiment with a *let-7b* expression plasmid (Nishino et al., 2008). EdU was added *in ovo* at time of electroporation to characterize the rate of pro-sensory cells proliferations. *Let-7b* overexpressing BPs harvested after 44 hours were severely malformed and growth retarded. To minimize potentially indirect effects on HC differentiation, we modified our experimental approach and harvested electroporated embryos after 18 hours at E5-E5.5 (HH26-HH27), coinciding with the onset of *Atoh1* expression. Consistent with *let-7’s* inhibitory role in cell proliferation, *let-7b* over-expression resulted in a significant decrease in EdU incorporation in SOX2(+) pro-sensory cells compared to left untreated controls (Fig. 4 A, B). However, *let-7b* overexpression failed to increase the number of *Atoh1*(+) cells. Instead, 12 out of the 14 analyzed specimens showed either a complete loss in *Atoh1* expression (n=6/14) or a reduction in the number of *Atoh1*(+) cells compared to control (n=6/14) (Fig. 4 C, D). Taken together these findings suggest that higher than normal *let-7* miRNA levels interfere with the induction of *Atoh1* and the subsequent differentiation of pro-sensory cells into HCs.

**Figure 4.**
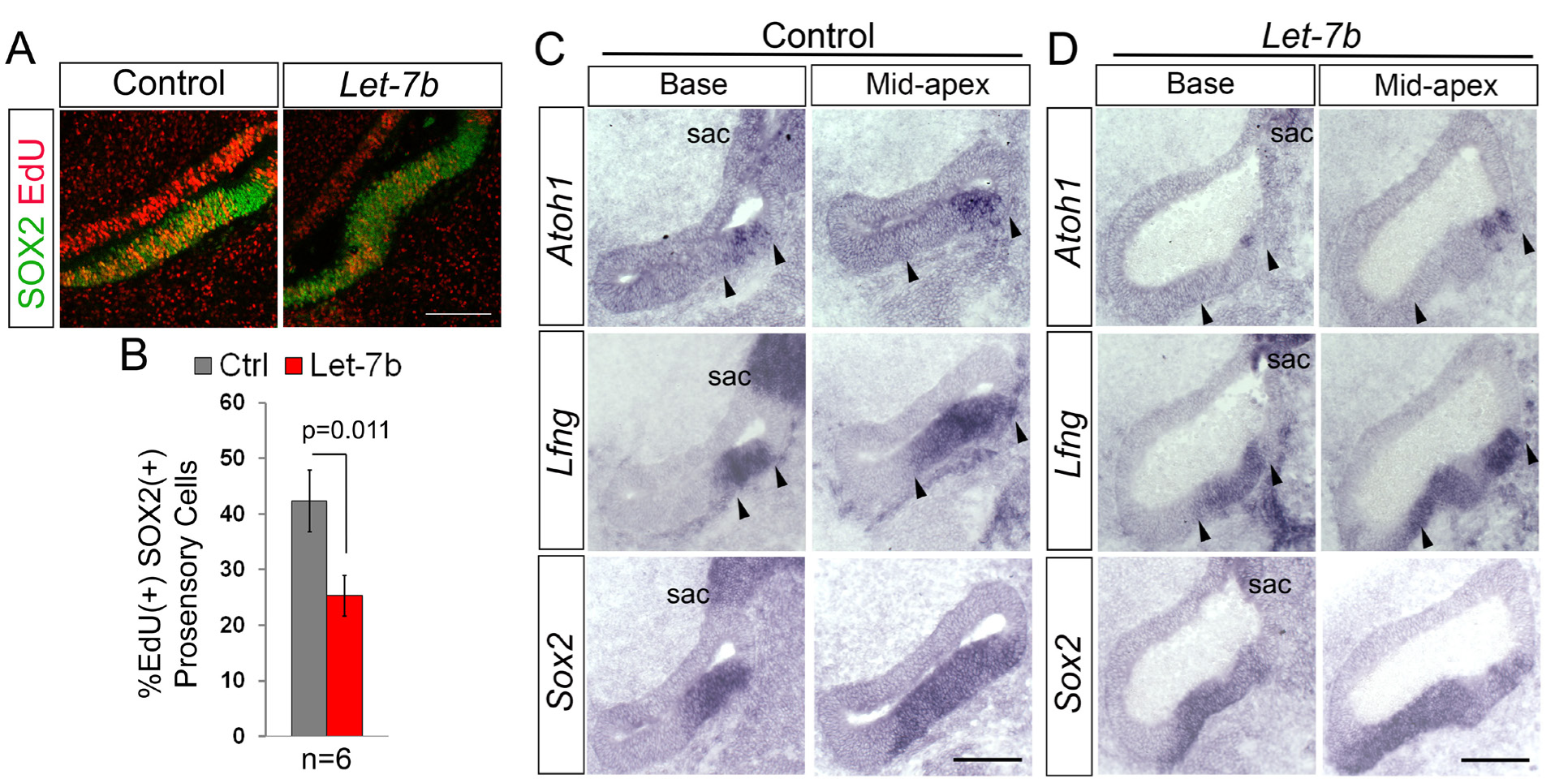
*Let-7* over-expression reduces pro-sensory cell proliferation and delays *Atoh1* induction in the developing BP. Shown are adjacent sections through the base or mid-apex of untreated control BPs (B-left panel, D) and *let-7b* electroporated BPs (B-right panel, E) 18 hours after electroporation and EdU treatment at E4.5. **(A-B) *Let-7b* overexpression reduces pro-sensory cell proliferation.** EdU incorporation in SOX2(+) pro-sensory cells is significantly decreased in the mid-apex of *let-7b*-electroporated BPs (A, right panel) compared to corresponding control BPs (A, left panel). (B) Quantification of A. Data are mean ± SEM (n = 6 animals per group, p=0.011 by 2-tailed Student’s t-test, p ≤ 0.05 is deemed significant). **(C-D) *Let-7b* overexpression delays HC differentiation**. The number of *Atoh1*(+) HC precursors/ HCs within the *Lfng*(+) *Sox2*(+) sensory epithelium is reduced in the *let-7b*-treated BP compared to control. Note, that the *let-7b*-treated BP is malformed. Abbreviation sac, sacculus. Scale bars 100 µm.

### *Let-7* miRNAs negatively regulate CHD7 expression in the developing chicken BP

Furthermore, we noted that in response to *let-7b* over-expression the size of the acoustic ganglion was severely reduced compared to control (Fig. 5 D, yellow asterisks; n=5/5). To identify *let-7* target genes involved in HC differentiation, we screened whether *Atoh1* or any of *Atoh1’s* upstream regulators EYA1, SIX1(Ahmed et al., 2012), SOX2 (Kempfle et al., 2016; Kiernan et al., 2005; Neves et al., 2012) and β-catenin (Ctnnb1) (Shi et al., 2010) contained *let-7-5p* binding sites. TargetScan 7.2. (Agarwal et al., 2015) and microT-CDS (Paraskevopoulou et al., 2013), two complementary miRNA prediction software packages, failed to uncover *let-7-5p* target sites within the mRNA sequence of chicken *Atoh1* or its upstream regulators. However, we identified a high probability *let-7-5p* binding site in the 3’ untranslated region (UTR) of chicken *Chd7* gene (Fig. 5 A). This binding site is evolutionary conserved and is present in the 3’ UTR sequence of the human and murine *Chd7* transcript (Fig. 5 B). The vertebrate *Chd7* (Chromo-Helicase-DNA binding protein 7) gene encodes for an ATP-dependent chromatin remodeling enzyme that amongst others is critical for the morphogenesis and neuro-sensory development of the mammalian inner ear (Hurd et al., 2010). Furthermore, CHD7 has been shown to cooperate with SOX2 in the transcriptional regulation of the Notch ligand JAGGED1 (JAG1) during early otic development (Engelen et al., 2011). JAG1-meditated Notch signaling is essential for the specification of auditory and vestibular pro-sensory cells in the developing inner ear (Daudet and Lewis, 2005; Kiernan et al., 2006; Pan et al., 2010). Interestingly, mice carrying heterozygous mutations in the *Jag1* gene largely lack the 3^rd^ row of outer hair cells (Kiernan et al., 2001; Tsai et al., 2001), somewhat resembling the cochlear phenotype reported for *let-7*g overexpressing mice (Golden et al., 2015).

**Figure 5.**
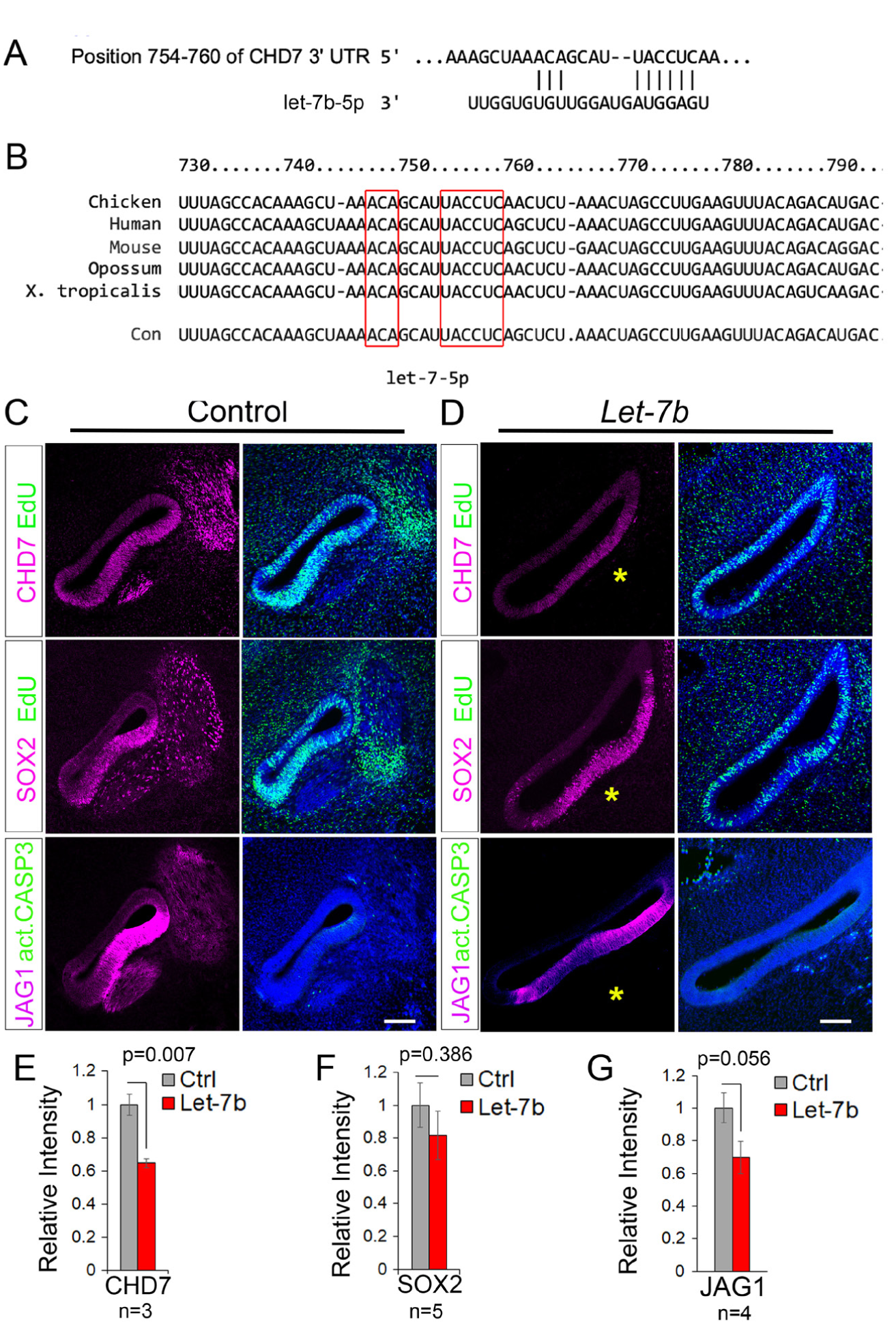
Let-7 miRNAs negatively regulate CHD7 expression in the developing BP. **(A-B) CHD7 is an evolutionary conserved let-7 target gene.** TargetScan Release 7.2 software was used to identify conserved *let-7-5p* binding sites in regulatory genes. The 3’UTR-sequence of the chicken Chd7 gene contains a high probability *let-7b-5p* binding site (A), which is highly conserved across vertebrate species (red boxes) (B). **(C-G) *Let-7b* overexpression reduces CHD7 and JAG1 expression in the developing BP**. Shown are confocal images of adjacent sections through the mid to mid-apex of untreated control BPs (C) and *let-7b* electroporated BPs (D) 18 hours after electroporation. SOX2 and JAG1 immuno-staining mark the pro-sensory domain. Activated Caspase3 (act-CASP3) immuno-staining labels apoptotic cells, phospho-Histone3 (p-H3) immuno-staining labels mitotic cells. Note, the auditory vestibular ganglion is severely reduced in size (yellow asterisk), and the BP is malformed in response to *let-7b* overexpression. (E-G) Relative intensity of CHD7 (E), SOX2 (F) and JAG1 (G) immuno-fluorescence in sections of control and *let-7b-*electroporated BPs. Data are mean ± SEM (n= number of animals per group, p-value calculated using 2-tailed Student’s t-test, p ≤ 0.05 deemed significant). Scale bars 100 µm.

We found that in the developing chicken BP (E4.5+18hrs), CHD7 protein was highly expressed in SOX2(+) JAG1(+) pro-sensory cells, as well as in flanking non-sensory epithelial cells. In addition, CHD7 was expressed in mitotic (EdU positive) and post-mitotic (EdU negative) cells within the developing acoustic ganglion (Fig. 5 C). As observed in our earlier experiments, pro-sensory SOX2 expression within the developing BP was not significantly altered by *let-7b* overexpression (Fig.5 C, D, F). However, consistent with being a target of *let-7* regulation, we found that in response to *let-7b* overexpression CHD7 protein expression was significantly reduced within the developing BP epithelia (Fig. 5 C, D, E). Furthermore, *let-7b* overexpression led to a mild reduction in JAG1 protein expression in pro-sensory cells (Fig.5 C, D, G). Taken together, these findings indicate that *let-7* miRNAs negatively regulate the expression of CHD7 in the developing BP, which in turn may impact the function and expression of SOX2 and JAG1 respectively.

### *Let-7* miRNAs repress CHD7 expression in the murine brain, retina and inner ear

To establish whether *let-7s* repressive function in *Chd7* gene expression is evolutionary conserved, we conducted *let-7* overexpression experiments in the developing mouse embryos using the well characterized *Rosa26*-*iLet-7g* mouse model. In this double transgenic mouse model a degradation-resistant form of *let-7g* is ubiquitously overexpressed throughout the body in the presence of doxycycline (dox) (Zhu et al., 2011). We administered dox to pregnant dams starting at stage E10.5 and harvested *let-7*g overexpressing embryos and their single transgenic control littermates for further analysis at stage E11.5 (Fig. 6) and E13.5 (Fig. 7). At stage E11.5, CHD7 protein is highly expressed within the developing otocyst and neuroblasts of the cochlear vestibular ganglion (CVG) (Hurd et al., 2010) (Fig. 6 A, control) as well the developing brain (Gage et al., 2015) (Fig. 6 D, control). By contrast, otic and brain-specific CHD7 expression was severely reduced in stage E11.5 *let-7g* overexpressing embryos (*iLet-7g*) (Fig. 6 A, D, *iLet-7g*). Consistent with previous reports that CHD7 positively regulates JAG1 expression in the developing otocyst (Engelen et al., 2011), the observed severe reduction in otic CHD7 expression was accompanied by a mild reduction in otic JAG1 expression (Fig. 6 A). Furthermore, consistent with *let-7’s* repressive role in cell proliferation, we found that the number of mitotic cells within the developing otocyst was significantly reduced in response to *let-7g* overexpression, as indicated by the significantly lower number of phopho-histone 3 (p-H3)(+) cells (Fig. 6 A, B). By contrast, *let7* overexpression did not alter the number of otic cells undergoing apoptosis, as indicated by the unchanged number of activate caspase 3 (CASP3)(+) cells (Fig. 6 A, C). Next, we analyzed the protein expression of CHD7 and TRIM71 (positive control) in whole brain lysates from E11.5 *iLet-7g* embryos and wild type littermates. (Fig. 6 E, F). Consistent with previous reports, our western blot-based analysis revealed that TRIM71 expression was severely reduced in the developing brain in response to *let-7g* overexpression (Lin et al., 2007). Furthermore, we found that *let-7g* overexpression causes a significant reduction in brain-specific CHD7 protein expression.

**Figure 6.**
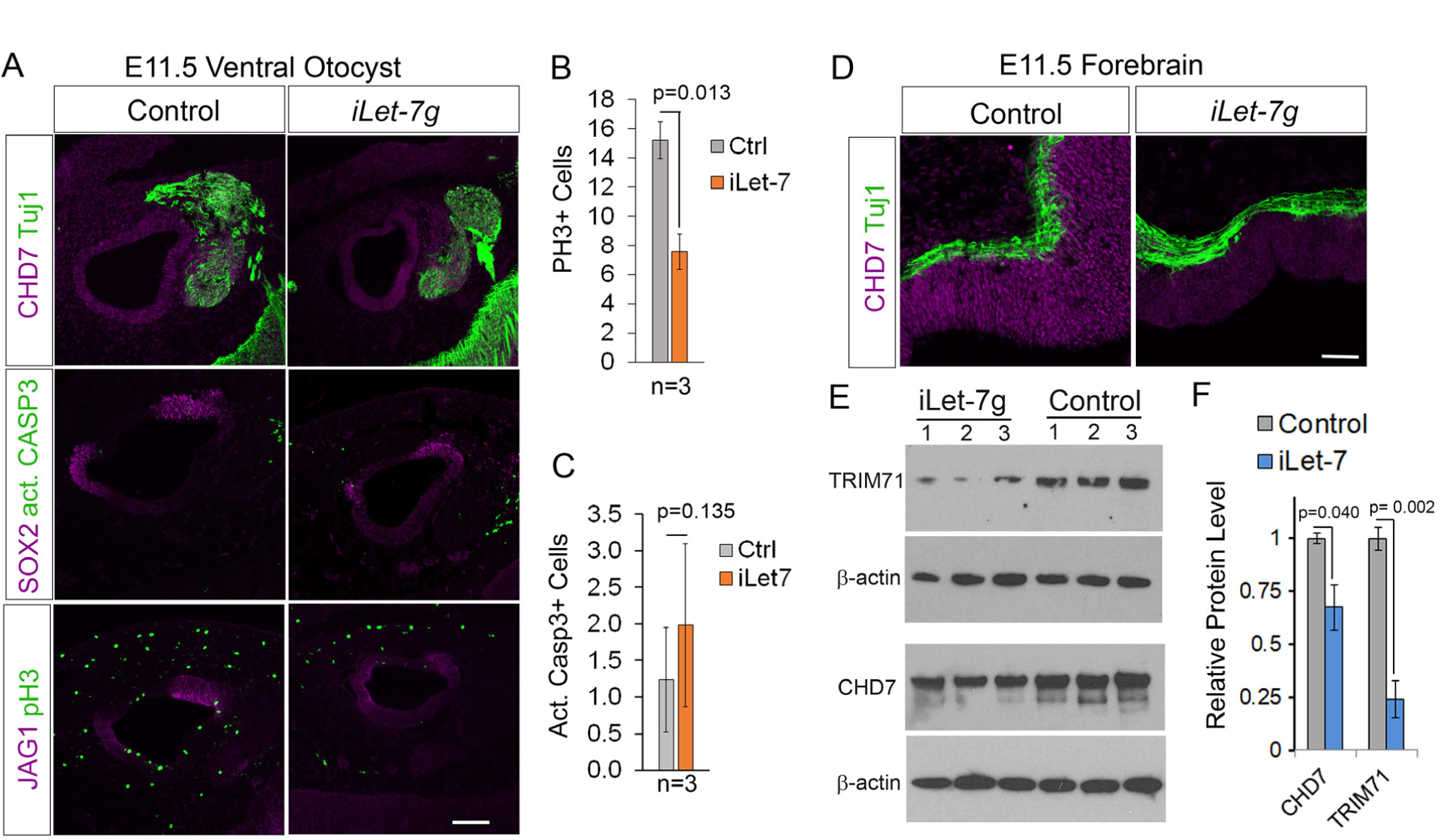
Let-7 miRNAs negatively regulate CHD7 expression in the murine otocyst and developing brain. We administered dox to pregnant dams starting at stage E10.5 and harvested *let-7g* overexpressing embryos and their single transgenic control littermates for further analysis at stage E11.5. **(A) *Let-7g* overexpression reduces otic CHD7 protein expression**. Shown are confocal images of adjacent sections through the ventral otocyst of control and *iLet-7g* littermates. Tuj1 immuno-staining marks neurons, SOX2 and JAG1 marks the pro-sensory domain. act-CASP3 labels apoptotic cells, p-H3 labels mitotic cells. **(B-C)** Quantification of mitotic pH3(+) cells (B) and apoptotic act. Casp3 (+) cells (C) in control (Ctrl, gray bar) and *let-7g* overexpressing (*iLet-7g*, orange bar) otocysts. Data expressed as mean ±SEM (n= 3 animals per group, p-value calculated using 2-tailed Student’s t-test, p ≤ 0.05 deemed significant). (**D-F) *Let-7g* overexpression significantly reduces CHD7 protein expression in the developing brain.** Shown are confocal images of CHD7 and Tuj1 immuno-stained sections through the forebrain of control and *iLet-7g* littermates. (F) Western blot analysis of CHD7 and TRIM71 expression in whole brain extracts of E11.5 *iLet-7* transgenic and control littermates. Beta actin was used as loading control. (F) Quantification of F. Graphed is the normalized TRIM71 and CHD7 protein levels within the brain of *iLet-7g* transgenic (blue bar) and control (grey bar) littermates. Data expressed as mean ±SEM (n= 3 animals per group, p-value calculated using 2-tailed Student’s t-test, p ≤ 0.05 is deemed significant). Scale bars 100 µm.

**Figure 7.**
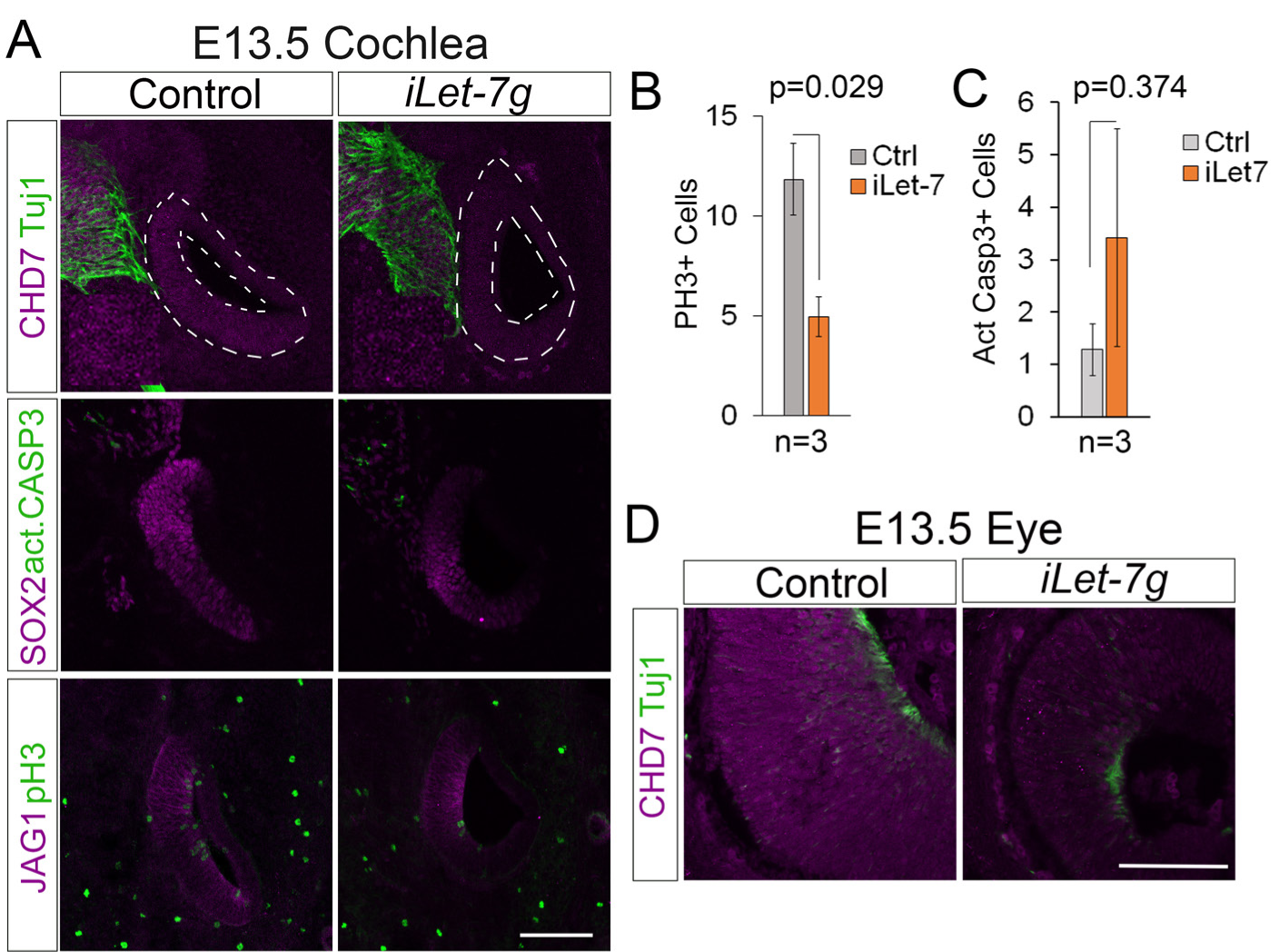
*Let-7* miRNAs negatively regulate CHD7 expression in the developing retina and cochlea. We administered dox to pregnant dams starting at stage E10.5 and harvested let-7g overexpressing embryos and their single transgenic control littermates for further analysis at stage E13.5. **(A) *Let-7g* overexpression reduces cochlear CHD7 protein expression**. Shown are confocal images of adjacent sections through the developing cochlear duct of control and *iLet-7g* littermates. Tuj1 immuno-staining marks neurons, SOX2 and JAG1 marks the cochlear pro-sensory domain. act-CASP3 labels apoptotic cells, p-H3 labels mitotic cells. **(B-C)** Quantification of mitotic pH3(+) cells (B) and apoptotic act. Casp3 (+) cells (C) in control (Ctrl, gray bar) and let-7g overexpressing (iLet-7g, orange bar) cochleae. Data expressed as mean ±SEM (p-value calculated using 2-tailed Student’s t-test, p ≤ 0.05 deemed significant). (**D) *Let-7g* overexpression reduces CHD7 protein expression in the developing retina.** Shown are confocal images of CHD7 and Tuj1 immuno-stained sections through the retina of control and *iLet-7g* littermates. Scale bars 100 µm.

At stage E13.5, CHD7 protein is highly expressed in the developing retina (Fig. 7 D, control) and in neural precursors of the cochlear spiral ganglion and at somewhat lower levels in cochlear pro-sensory cells and flanking non-sensory epithelial cells (Fig. 7 A, control) (Hurd et al., 2007). By contrast, retinal CHD7 expression (Fig. 7 D, *iLet-7g*), as well as cochlear CHD7 expression were severely reduced in E13.5 *iLet-7g* transgenic embryos. Interestingly, no reduction in JAG1 protein expression was observed in *let-7g* overexpressing cochlear tissue compared to control, indicating that CHD7 function may not be required for JAG1 expression at this later stage. However, we noticed that the size of the cochlear duct and the size of its SOX2(+) pro-sensory domain were reduced in *let-7g* overexpressing embryos compared to control (Fig. 7 A, *iLet-7g*). This was unlikely due to an increase in cell death, as both control and *let-7g* overexpressing cochlear epithelia contained only few scattered act-CASP3(+) cells (Fig. 7 A, C). However, *let-7g* overexpression significantly reduced the number of mitotic pH3 (+) cells within the developing cochlear duct (Fig. 7 A, B), suggesting that premature cell cycle exit may have contributed, to the smaller cochlea and the smaller pro-sensory domain observed in *let-7g* overexpressing embryos.

Taken together, our analysis establishes *let-7* miRNAs as negative regulators of CHD7 protein expression in the developing inner ear, brain and retina. CHD7 has dosage specific functions and the here described regulation of CHD7 by *let-7* miRNAs may be critical to fine tune CHD7 protein levels during sensory and neuronal development.

## DISCUSSION

The highly specialized mammalian and avian auditory sensory organs have evolved independently (Fritzsch et al., 2013). Yet, many of the regulatory gene networks that control patterning, specification and differentiation of the auditory sensory epithelium appear to be conserved among avians and mammals (Munnamalai and Fekete, 2013). In the mammalian cochlea cell cycle exit and differentiation of pro-sensory cells is initiated at opposing ends of the developing cochlear duct. Pro-sensory cells located at the cochlear apex are the first to exit the cell cycle but are the last to differentiate (Chen et al., 2002). Spatial segregation between pro-sensory cell cycle withdrawal and differentiation, even though less pronounced, has been also reported for the developing chicken BP (Cotanche and Sulik, 1984; Katayama and Corwin, 1989). Indeed, our analysis revealed that in the developing chicken BP terminal mitosis of pro-sensory cells occurs first in the proximal (basal) half of the BP, whereas HC differentiation initiates in the distal (apical) half of the BP.

What times and coordinates these seemingly uncoupled events? We have recently shown that the LIN28B/*let-7* pathway is active in the developing murine auditory sensory organ and plays a key role in timing pro-sensory cell cycle exit and differentiation (Golden et al., 2015). Expanding on these recent findings, we now show that *let-7* miRNAs are essential for the proper timing of terminal mitosis and differentiation the developing chicken auditory organ. In particular, we find that expression of mature *let-7* miRNAs sharply rises around the time of terminal mitosis and using a RNA sponge to sequester functional *let-7* miRNAs, we show that disruption of *let-7* function in pro-sensory cells delays their onset of terminal mitosis and differentiation, whereas overexpression of the *let-7* miRNA species *let-7b* accelerated terminal mitosis. How do *let-7* miRNAs exert control over pro-sensory cell proliferation and maintenance? We show that the evolutionary conserved *let-7* target genes *Hmga2, Lin28b* and *Trim71* are highly expressed in undifferentiated BP (E5.5), but are near absent in the differentiated BP. The RBPs TRIM71 and LIN28B and the chromatin associated protein HMGA2 are key regulators of stemness and have been shown to promote stem cell/ progenitor cell proliferation in both neuronal and sensory lineages (Maller Schulman et al., 2008; Nguyen et al., 2017) #334;Yang, 2015 #3185;Nishino, 2008 #3186; Parameswaran, 2014 #3293}. Thus, it is likely that *let-7* miRNAs time auditory pro-sensory cell cycle exit in the developing chicken BP, at least in part, through repressing *Hmga2, Lin28b* and *Trim71* expression. Support for such model comes from our recent study that revealed that prolonged LIN28B expression maintains auditory pro-sensory cells in a proliferative undifferentiated state (Golden et al., 2015).

The role of *let-7* miRNAs in the differentiation of neural-sensory progenitors is complex and highly dependent on the cellular context. For instance, overexpression of *let-7b* stimulates the differentiation of cultured retinal progenitors and neuronal stem cells through targeting nuclear receptor TLX (NR2E1) (Ni et al., 2014; Zhao et al., 2010). By contrast, *let-7b* or *let-7i* overexpression in human ESC derived neuronal progenitor cells inhibits neuronal differentiation, as it reduces the expression of the pro-neural gene *ASCL1* (*MASH1*) (Cimadamore et al., 2013). We previously reported that overexpression of *let-7g* interferes with auditory HC differentiation in mice. Expanding on theses previous findings, we provide evidence that *let-7* levels have to be tightly controlled during auditory sensory differentiation. We show that in the developing chicken BP higher than normal as well as lower than normal levels of *let-7* miRNA expression significantly delay auditory HC differentiation. This seemingly paradoxical finding suggest that *let-7* miRNAs, in addition to targeting genes critical for pro-sensory cell self-renewal, target genes involved in HCs differentiation. Auditory HC formation depends on the function of the transcription factor ATOH1 (Bermingham et al., 1999) and its upstream regulators EYA1, SIX1(Ahmed et al., 2012), SOX2 (Kempfle et al., 2016; Neves et al., 2012) and β-catenin (Shi et al., 2010). We failed to identify *let-7-5p* binding sites within the mRNA sequence of these key regulatory genes. However, we uncovered a highly predictive *let-7-5p* binding site within the 3’UTR of the chicken, human and murine *Chd7* gene. The predicted interaction of *let-7-5p* miRNAs with the human CHD7 mRNA have been recently validated through genome wide interrogation of miRNA-target RNA duplexes (Helwak et al., 2013) and consistent with being a canonical target of *let-7* repression, we find that overexpression of *let-7b* or *let-7g* reduced the expression of CHD7 in the chicken or murine auditory organ respectively.

Heterozygous mutations in the human *CHD7* gene are a leading cause for CHARGE syndrome, a disorder that causes multiple birth defects including ocular coloboma, mental retardation, ear abnormalities and deafness (Pagon et al., 1981). Inner ear-specific ablation of the murine *Chd7* gene causes severe inner ear malformation, including severe cochlear hypoplasia and defects in neurogenesis (Hurd et al., 2010). The role of CHD7 in hair cell development has not yet been established. CHD7, an ATP-dependent chromatin remodeler, is preferentially found at active enhancer/promoter regions, and has been shown to control chromatin accessibility and transcriptional activation/repression in a cell lineage specific manner (Bajpai et al., 2010; Chai et al., 2018; Feng et al., 2013; Schnetz et al., 2009; Schnetz et al., 2010). In Drosophila, the CHD7 orthologue Kismet is required for atonal transcription in the developing eye (Melicharek et al., 2008). To date there is no evidence that CHD7 directly activates *Atoh1* transcription in vertebrates. However, there are several candidate genes through which CHD7 may impact the HC differentiation. CHD7 has been shown to physically interact with SOX2 and co-regulate a set of common target genes (Doi et al., 2017; Engelen et al., 2011). Thus, *let-7* induced reduction in CHD7 expression in the developing chicken and murine auditory organ may compromise the ability of SOX2 to activate *Atoh1* expression and subsequent HC formation (Kempfle et al., 2016; Neves et al., 2012). Furthermore, CHD7 has been recently shown to be required for the transcriptional activation of *Sox11* and *Sox4* in adult neuronal stem cells (Feng et al., 2013). The transcription factors SOX11 and SOX4 are broadly expressed in cochlear and vestibular pro-sensory cells and in their absence both auditory and vestibular HCs fail to form (Gnedeva and Hudspeth, 2015). Future investigations are warranted to address the role of CHD7 in the timing of *Atoh1* induction and HC differentiation.

## ACKNOLEDGMENTS

We thank the members of the A.D. Laboratory for the help and advice provided throughout the course of this study. We thank Dr. Doris K. Wu for *in situ* probes and Dr. Georg Dailey for *iLet-7* mice. The work was supported by NIDCD Grants DC000023 (L.E), DC011571 (A.D.), DC005211 (Sensory Mechanisms Research Core Center) and David M. Rubenstein Fund for Hearing Research (A.D.).

