## supplemental data for "*Let-7* miRNAs control auditory sensory progenitor behavior in the vertebrate inner ear"

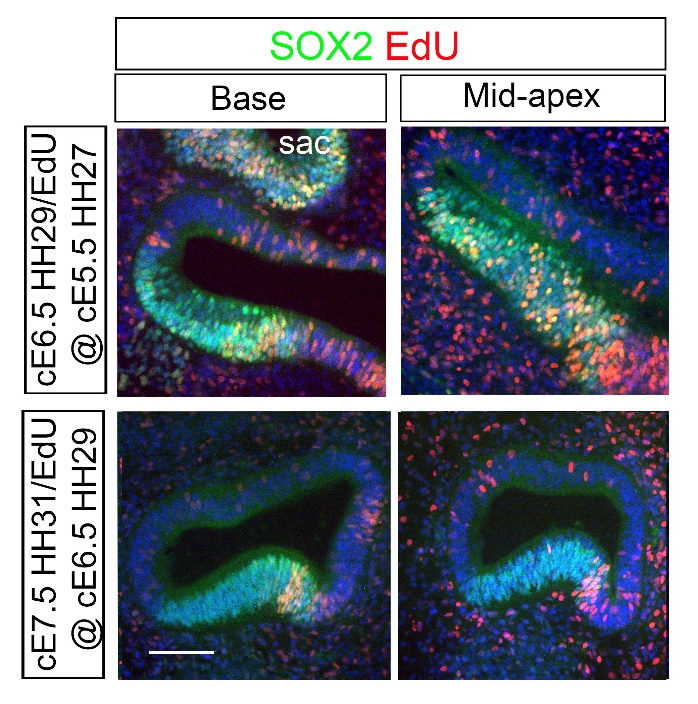

**Fig.S1. Cell proliferation ceases around stage E6.5 (HH29) in the developing BP.** Tissue-sections through the wild-type BP base or mid-apex at E6.5 and E7.5 with addition of EdU at E5.5 and E6.5, respectively; SOX2 immuno-staining labels pro-sensory domain/ sensory domain. Abbreviation: sac, sacculus. Scale bar 100 µm.

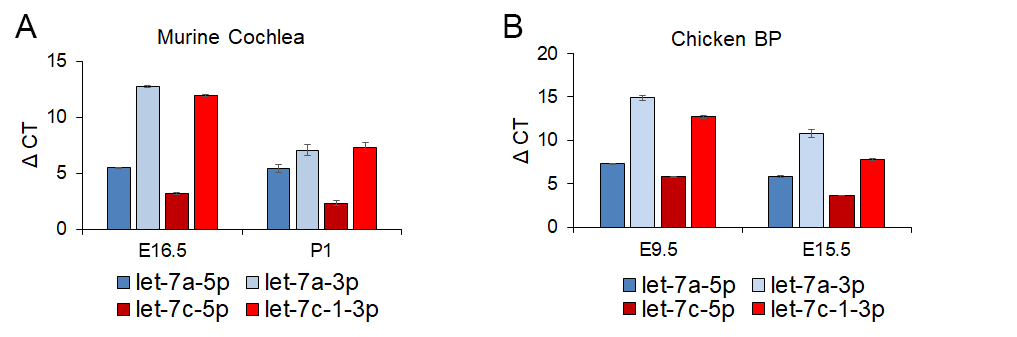

**Fig.S2: Let-7 passenger strand expression in the developing mammalian and avian auditory organ.** **(A-B)** *Let-7* lead strands *(let-7a-5p,* *let-7c-* *5p*) are expressed a much higher level (lower ΔCT) than their corresponding passenger strands *(let-7a-3p, let-7c-1-3p)* in the differentiating (E16.5) and terminal differentiated (P1) murine cochlea (A) as well as the differentiating (E9.5) and terminal differentiated (E15.5) chicken BP. Plotted on the y-axis are the difference in cycle threshold (ΔCT) between target genes and the reference gene U6. Note that ΔCT level of 12 or higher indicates that the transcript is expressed at a very low level. Data are mean ± SEM (n = 3 technical replicates).

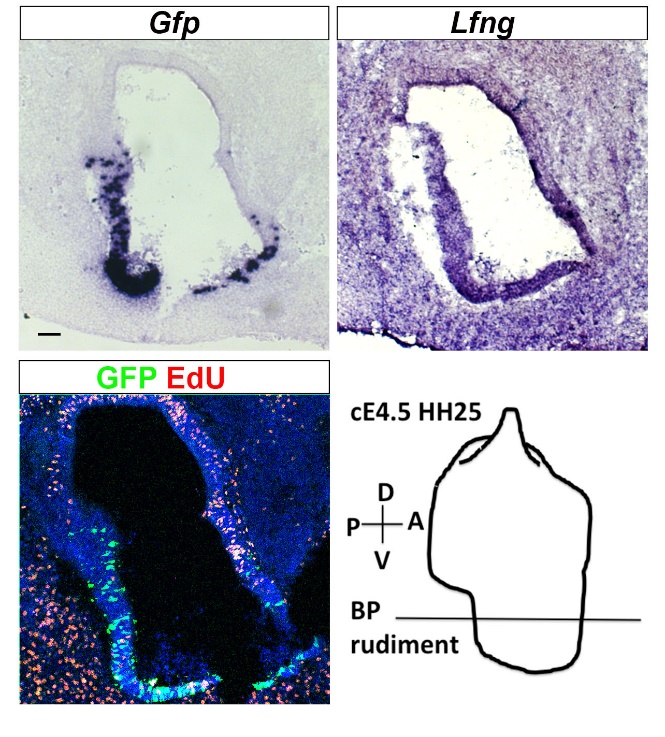

**Fig.S3. Gene transfer targeting the BP rudiment by in ovo electroporation.** Chicken embryos were electroporated at E4.5 (HH25) targeting GFP control expression plasmid to the ventral portion of the developing otocyst (see schematic). *Gfp* mRNA and GFP protein expression was analyzed in the adjacent tissue sections of the BP rudiment 6 hours later. Note *Lfng* mRNA expression marks pro-sensory cells within the developing BP. Scale bar 100 µm.

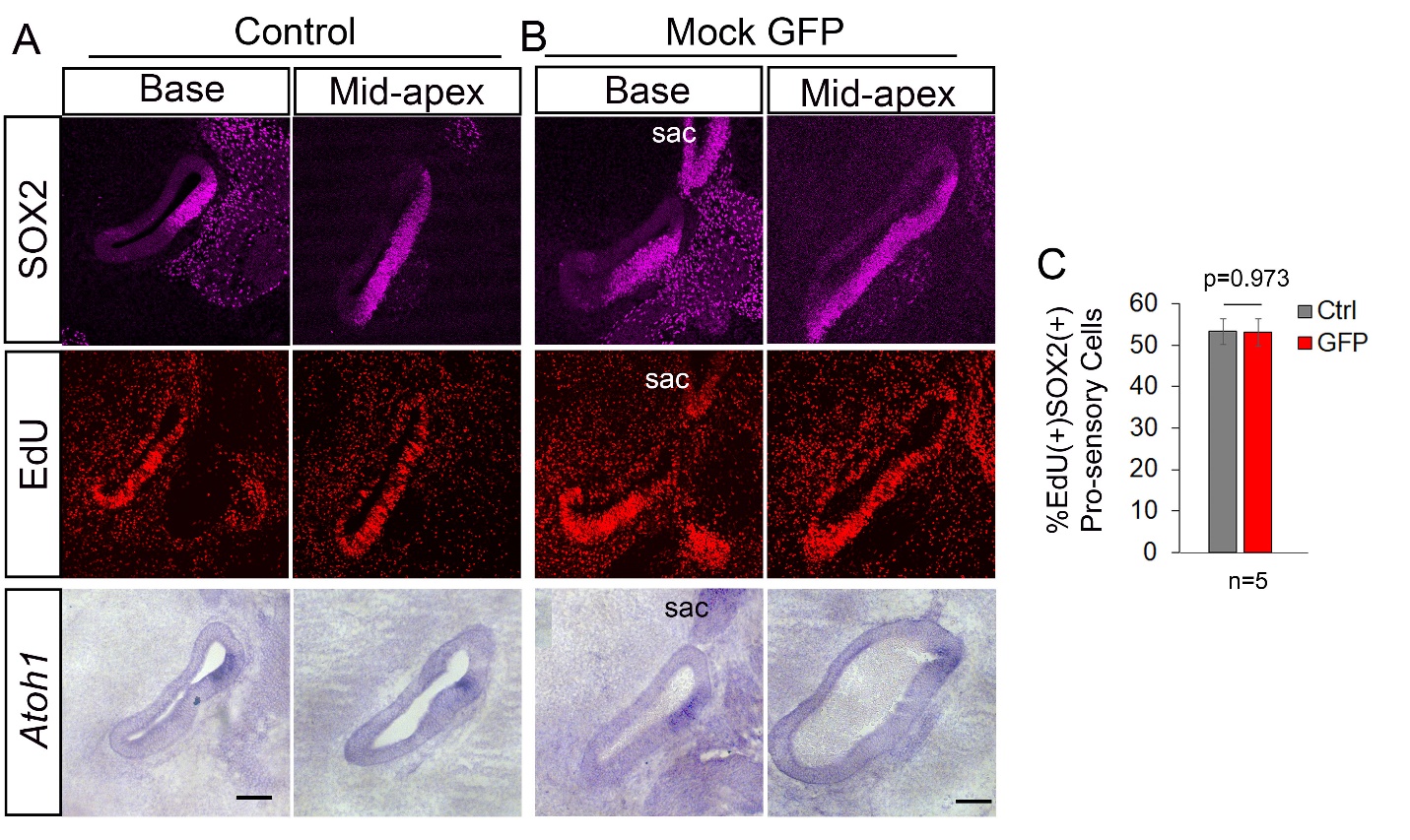

**Fig.S4 Mock electroporation of the developing BP with GFP does not alter the timing of terminal mitosis and HC differentiation.** (A-B) Shown are adjacent sections through the base or mid-apex of untreated control BPs (A) and mock *GFP* electroporated BPs (B) 18 hours after electroporation and EdU addition at E4.5. Note mock-electroporated BPs with incorporated EdU in SOX2(+) pro-sensory cells at a similar rate than control. Also, HC differentiation in mock electroporated BPs, as indicated by the number of *Atoh1*(+) HC precursors/ HCs, remained unchanged compared to control. (C) Graphed is EdU incorporation in SOX2(+) pro-sensory cells in control (gray bar) and mock-GFP electroporated BPs (red bar). Data expressed as mean ±SEM (n=5 animals per group, p=0.973 by 2-tailed Student’s t-test, p ˃0.05 is not significant). Scale bars 100 µm
